# Sequence characterization of the complete chloroplast genome of *Eupatorium fortunei*

**DOI:** 10.1101/2022.04.18.488663

**Authors:** Kan Yan, Juan Ran, Songming Bao, Nai Zhang, Wei Zhao, Yanni Ma, Marie Angela Ishimwe

**Author notes:** Corresponding author, Also the first author. These authors contributed equally to this work (Kan Yan, Juan Ran).

## Abstract

*Eupatorium fortunei* has been utilized as herbal medicine to cure various diseases in various regions of the world especially China, Korea, and other Asian countries. The available literature shows *Eupatorium fortunei* contain anti-metastatic, anti-angiogenic, anti-bacterial, and anti-oxidant, along with compounds to cure human cancer. In present study, *Eupatorium fortunei* was used as an experimental material, and its genome was sequenced based on high-throughput sequencing technology. We assembled the complete chloroplast genome of *Eupatorium fortunei* and obtained NCBI accession number (OK545755). We analyzed chloroplast genome structure of *Eupatorium fortunei* and systematic analysis was conducted, including the size of the entire genome sequence, genome volume, large single-copy (LSC), small single-copy (SSC), inverted repeat region (IR)-LSC/SSC boundaries, etc. The obtained results showed that chloroplast genome of *Eupatorium fortunei* is a typical tetrad structure with a total length of 152,401 bp and a sum of 133 annotated genes, including eight ribosomal RNA (rRNA)-encoding genes, 37 transfer RNA (tRNA)-encoding genes, 87 protein-encoding genes, and one pseudo-gene. A sum of 29 Single Sequence Repeats (SSR)s were identified for mono-nucleotide, di-nucleotide and tri-nucleotide microsatellites with mono-nucleotides in majority. The phylogenetic analysis revealed that *Eupatorium fortunei* is more closely related to *Litothamnus nitidus* and belongs to the same branch. The genome assembly and analysis of the chloroplast genome of *Eupatorium fortunei* provides more accurate evidence for further taxonomic identification and localization of *Asteraceae/Compositae* plant.

## Introduction

*Asteraceae* is the second-largest family of plant kingdom, having complex taxon in composition and largest *eudicots*, with 13 subfamilies, 1,689 genera, and 32,913 species(1). The *Asteraceae* family is distributed among all continents of the world, except Antarctica. *Eupatorium fortunei* is a genus of *Asteraceae* family, it consist of approximately 36 to 60 flowering plants species (2). *Eupatorium fortunei* is a perennial herb, growing in roadside, thickets and of ravines. The origin of *Eupatorium fortunei* is distributed in Shandong, Jiangsu, Hubei, Hunan, Yunnan, Sichuan, and other provinces in China, and a small amount in Japan, Korea, and other countries. It mainly contains coumarin, volatile oil, o-coumaric acid, triterpenoids, and muscadine hydroquinone. The available literature have shown that *Eupatorium fortunei* has a long history of medicinal use andas a whole herb to benefit dampness, strengthen the stomach, and clear summer heat. According to Chinese medicine, *Eupatorium fortunei* has pharmacological effects such as resolving dampness, relieving summer heat, awakening the spleen and appetite, anti-tumor, and anti-inflammatory and treating dampness, inflammation, and immune regulation influenza in the body. Meanwhile, *Eupatorium fortunei* plant has dense leaves, white or reddish flowers, pleasant fragrance, and high ornamental value.

Owing to *Eupatorium fortunei* medicinal importance it’s noteworthy to understand the genetic architecture of attributes for potential medical use. The recent development in high-throughput sequencing technology it has become convenient to study the chloroplast genome of both model and non-model plants (4). The chloroplast is an unique organelle, main site of photosynthesis and efficiently convert sunlight into energy in higher plants and in some algae, respectively. The chloroplast had its own genome ranged from 120-180 kn in higher plants (5, 6). Furthermore, it has lower molecular weight and numerous copies (7). The chloroplast genome is circular having single-copy region (SSC), a large single-copy region (LSC), and a pair of inverted regions (IRs) (8, 9). The first chloroplast genome were reported in liverwort followed by tobacco(11).

The rapid development in next-generation sequencing (NGS) technologies i.e., Illumina GenomeAnalyzer and Roche/454 GS FLX has made the sequencing of chloroplast genome both efficient and economical. So far, 4,100 plant’s chloroplast genome have been published and available at public repository e.g., NCBI database. The chloroplast genome also contain numbers of functional genes which hold potential to be utilized for species identification and evolutionary studies, widely adopted and accepted by researchers around the globe (12, 13). Moreover, chloroplast genome not only determine previously reported phylogeny but also increase the accuracy of phylogenetic trees. To comprehensively explore the interspecific relationships within the genus *Eupatorium*, the evolutionary relationships of *Eupatorium fortunei* is investigated, moreover, the taking into previous studies on *Asteraceae* are foundation knowledge to conduct present study. To investigate the phylogenetic position and genetic background of the *Eupatorium fortunei*, we sequenced the DNA of the *Eupatorium fortunei* specie. Furthermore, the complete chloroplast genome sequence of *Eupatorium fortunei* was assembled, analyzed and compared with the chloroplast genome of *Asteraceae* to explore its phylogenetic relationships and provide new insights into the taxonomy and systematics of *Eupatorium*.

## Materials and Methods

### Plant material and DNA extraction

Fresh leaves of *Eupatorium fortunei* were collected from Yangmingshan National Forest Park, Shuangpai County, Yongzhou City, Hunan Province, China (26°04’31.8 “N,111°55’33.8 “E) during May 2021. The fresh leaves were immersed in liquid nitrogen and later ground to fine powder using mortar and pestle, and genomic DNA was extracted using the Plant Genome Rapid Extraction Kit, and then DNA concentration and quality were determined using a NanoDrop 2000 Ultra Micro UV Spectrophotometer.

### DNA sequencing and sequence assembly

DNA sequencing was performed at Wuhan Bena Technology Services Co. The extracted high-quality DNA was used for 350 bp shotgun library construction, and high-throughput sequencing was performed on the Illumina NovaSeq 6000 sequencing platform. The raw image data files obtained from sequencing were transformed into raw data stored in FASTQ file format (Raw reads) by Base Calling analysis. The raw data were then filtered for low-quality sequences, splice sequences, etc. using Burrows-Wheeler Aligner’s sequence comparison tool and Samtools toolkit to obtain clean reads, to ensure the reliability of results were stored in FASTQ format for subsequent analysis, public database submission and publication.

Pre-splicing of the genome was performed using the NOVOPlasty software, splicing results were Blastn and compared on NCBI (https://www.ncbi.nlm.nih.gov/) database to select reference sequences for subsequent genome assembly. The collinearity analysis was undertaken on the pre-spliced results files using the nucmer command in Mummer to determine reference sequence relative positions and orientations in the genome to continue the construction of the cp genome sequence. Moreover, results were verified whether they are joined into a loop to obtain the complete genome sequence.

### Chloroplast genome annotation

Using the chloroplast annotation tool GeSeq (https://chlorobox.mpimp-golm.mpg.de/geseq.html), uploaded the genome sequence and reference genome GenBank file, and selected the parameters to obtain the preliminary annotation result file. The preliminary annotation files of *Eupatorium fortunei* chloroplast genome were manually corrected with the help of software Notepad++ and software Geneious 8.0.4, for each gene where necessary. The annotation files of chloroplast genome sequence were uploaded to OrganellarGenomeDRAW (OGDRAW, http://ogdraw.mpimp-golm.mpg.de) with default setting to determine the order of gene alignment and the position of the inverted repeats (IRs) with the large single-copy (LSC) region and the small single-copy (SSC) region, and finally to generate the physical mapping of the cyclic chloroplast genome of *Eupatorium fortunei*. The chloroplast genome sequence and annotation files of *Eupatorium fortunei* were submitted to the GenBank database to obtain the genome registration number.

### Simple sequence repeat analysis

The microsatellite identification tool (MISA, https://webblast.ipk-gatersleben.de/misa/) was used to identify and localize potential Single Sequence Repeats (SSR) sites in the complete cp genome sequence. Parameters setting: the minimum repeat numbers were set as 10 repeat units for mono-nucleotides, and five for di-, tri-, tetra-, penta-, and hexa-nucleotides whereas, the maximum sequence length between two composite SSRs was set 20 bp.

### Comparative analysis of chloroplast genomes

To understand the characteristics and differences, the chloroplast genomes of *Eupatorium fortune* were compared with those of nine other *Asteraceae* plants species based on the annotated information of chloroplasts. We used IRScope software (https://irScope.shinyapps.io/Irapp/) to generate a comparison diagram of the inverted repeat region (IR) boundary, to quantify the gene and neighboring gene characteristics at each boundary point (LSC-IRa, IRa-SSC, SSC-IRb, IRb-LSC). The distribution of genes in the boundary regions was then compared to shown their differences, demonstrating the diversity of chloroplast genome structures in different *Asteraceae* plant species.

There are differences in the organization of plant chloroplasts and having numbers of internal mutations. By using the online software mVISTA (https://genome.lbl.gov/vista/mvista/submit.shtml), the *Asteraceae* cp genome sequences was performed using *Eupatorium fortunei* as the reference genome, which can visualize the overall sequence similarity and variation in hotspot regions. Then Shuffle-LAGAN global alignment mode was selected, and other parameters were set at default values to find gene re-arrangements and inversions.

### Phylogenetic Analyses

Complete chloroplast genome sequences of 12 *Asteraceae* species were selected from NCBI (https://www.ncbi.nlm.nih.gov/) for cluster analysis to infer phylogenetic relationships, including nine *Mikania*, one *Stevia*, one *Ageratina*, and one *Litothamnus* plant, respectively. Multiple sequence alignment of nucleotide sequences was performed using MAFFT v7.308, and the Neighbor-Joining (NJ) method in MEGA7 was used to cluster 13 cp genomic sequences including *Eupatorium fortunei*. The bootstrap method was used to construct a phylogenetic tree using the bootstrap value test with 1000 replications.

## Results

### Sequencing and assembly of the chloroplast genome of *Eupatorium fortunei*

The *Eupatorium fortunei* chloroplast genome coverage was 100% using high-throughput sequencing platform Illumina HiSeq 6000, and the raw short sequence data (Sequenced Reads) of the double-ended Illumina reads obtained from sequencing were approximately 9.4 GB, which contained information on the bases of the sequenced (Reads) and its corresponding sequencing quality information. to ensure the quality of information analysis, the reads with connectors and low quality were filtered, and after data filtering, the clean data volume was 320M reads, which were utilized for subsequent information analysis. After preliminary splicing, three valid Contigs were obtained, contig 01 with 133,092bp, contig 02 with 19,561bp, and contig 03 with 19,309bp. The complete cp genome of *Eupatorium fortunei* was obtained after genome fragment splicing and gap-filling.

### Basic structural features of the chloroplast genome

The total length of the *Eupatorium fortunei* chloroplast genome is 152,401 bp (Fig 1). The cp genome contains four characteristic regions: LSC region of 83,032 bp, SSC region of 19,309 bp, and a pair of IRs (IRA and IRB) of 25,030 bp in length. Analysis of the base composition of the complete cp genome sequence revealed a sum of 31.1% consisting, adenine 18.5%, cytosine 19.1%, guanineand 31.3% for thymine. The overall GC content were 37.6%, very close to other *Asteraceae* species. In addition, the GC content were unevenly distributed among the regions of the cp genome, high GC content in the IR region, accounting for 43.06%, and a relatively low GC content in the LSC and SSC regions, 35.71% and 31.48%, respectively.

**Fig 1.**
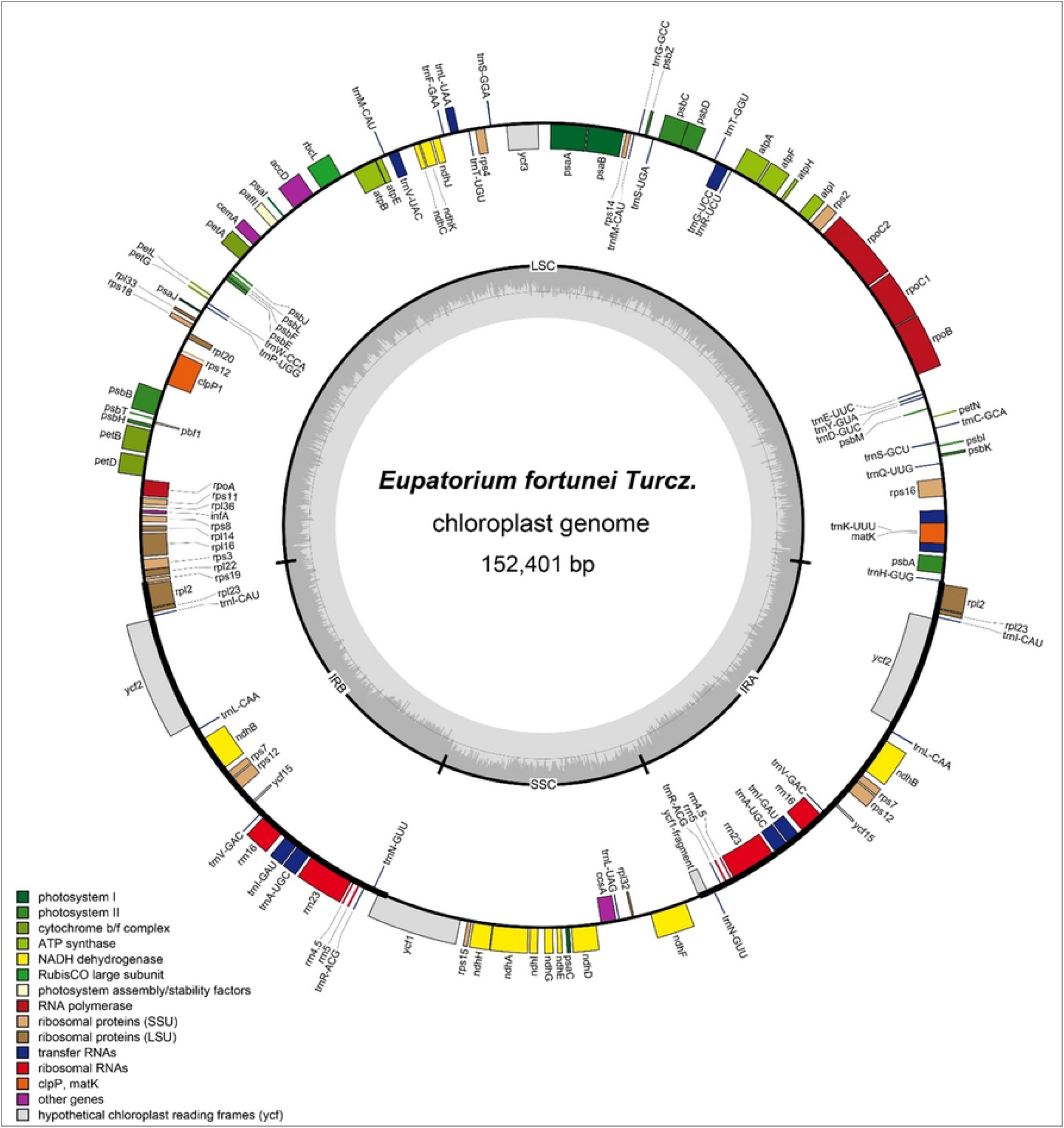
The chloroplast genome of *Eupatorium fortunei*. From the center going outward, the four circles indicate scattered forward and reverse repeats, tandem repeats, microsatellite sequences identified, and gene structure of the plastome.

The *Eupatorium fortunei* cp genome consists of 133 genes, including 87 protein-coding genes (PCGs), 08 rRNA-coding genes and 37 tRNA-coding genes, and one pseudo-gene. In *Magnoliophyta*, the structure and sequence composition of the chloroplast genome are highly conserved. The gene composition of *Eupatorium fortunei* is the same as that of most *Magnoliophyta* plant chloroplast genomes. No major gene gain or gene loss was found, and it has the typical structure of the *Magnoliophyta* plant chloroplast genome. (26). Among them, *Eupatorium fortunei* chloroplast encodes 115 single-copy, and 18 double copies genes, including seven PCGs (*rpl23, rpl2, rps7, rps12, ndhB, ycf2, ycf15*), seven tRNA genes (*trnA-UGC, trnI-CAU, trnL-CAA, trnI-GAU*, *trnN-GUU*, *trnR-ACG* and *trnV-GAC*) genes and four rRNA genes were also present in the IR region in two copies. Based on function chloroplast genes can be classified into three categories: category 1 include 74 genes related to transcription and translation; category 2 contained 45 genes related to photosynthesis, and category 3 includes 14 genes related to biosynthesis of substances such as amino acids and fatty acids and containing some genes of unknown function (Table 1).

**Table 1.**
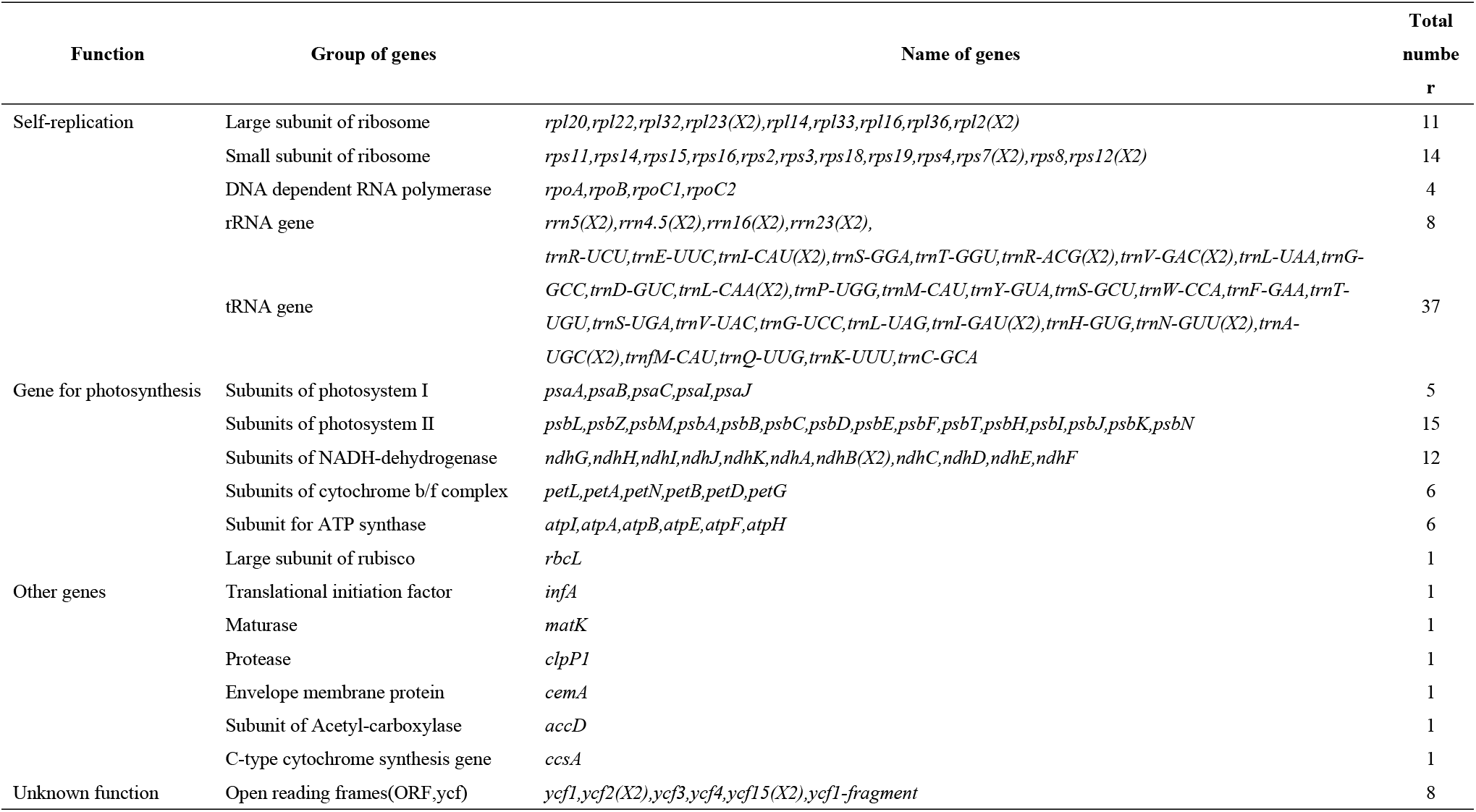
List of genes in the chloroplast genome of *Eupatorium fortunei*.

### SSR analysis

Simple sequence repeats (SSRs) are efficient molecular markers with advantage of being abundant, high reproducibility, and co-dominant inheritance, as well as, uniparental inheritance, relative conservation, which makes them best fit for for species identification and analysis of genetic differences both at population and individual levels (27).

A sum of 29 SSRs were recognized as potential cp markers in the chloroplast genome of *Eupatorium fortunei*, and SSRs were disproportionately spaced at cp genome. The SSRs frequency of the LSC region was significantly higher than that of the IR and SSC regions, as shown in Table 2. Among the SSRs, 26 mononucleotide, 2 dinucleotide and 1 trinucleotide microsatellites were identified. The mononucleotide microsatellites accounted for 89.67% of the SSRs in the *Eupatorium fortunei* cp genome. All mononucleotide and two dinucleotide microsatellites were composed of A and T except one mononucleotide microsatellite composed of C. Besides, most of the SSRs were distributed in LSC (68.97%), followed by SSC (24.14%), and IRs accounted for less than 7%.

**Table 2.** Repeat sequences and their distribution within *Eupatorium fortunei* chloroplast genome.

| Number | SSR type | Motif | Size | Start | End |
| --- | --- | --- | --- | --- | --- |
| 1 | p1 | (T)10 | 10 | 2299 | 2308 |
| 2 | p1 | (C)12 | 12 | 5418 | 5429 |
| 3 | p1 | (T)10 | 10 | 9543 | 9552 |
| 4 | p1 | (A)10 | 10 | 13302 | 13311 |
| 5 | p1 | (T)12 | 12 | 16571 | 16582 |
| 6 | p1 | (A)10 | 10 | 18356 | 18365 |
| 7 | p1 | (T)13 | 13 | 24915 | 24927 |
| 8 | p2 | (TA)6 | 12 | 26593 | 26604 |
| 9 | p1 | (T)14 | 14 | 27969 | 27982 |
| 10 | p1 | (T)10 | 10 | 30842 | 30851 |
| 11 | p2 | (AT)7 | 14 | 43871 | 43884 |
| 12 | p1 | (T)14 | 14 | 46306 | 46319 |
| 13 | p1 | (T)12 | 12 | 53844 | 53855 |
| 14 | p1 | (T)11 | 11 | 58539 | 58549 |
| 15 | p1 | (T)10 | 10 | 69445 | 69454 |
| 16 | p1 | (T)10 | 10 | 69481 | 69490 |
| 17 | p1 | (T)10 | 10 | 70461 | 70470 |
| 18 | p1 | (T)10 | 10 | 70625 | 70634 |
| 19 | p1 | (T)10 | 10 | 77060 | 77069 |
| 20 | p1 | (T)10 | 10 | 79989 | 79998 |
| 21 | p1 | (A)11 | 11 | 106342 | 106352 |
| 22 | p3 | (GAA)5 | 15 | 108410 | 108424 |
| 23 | p1 | (T)12 | 12 | 108521 | 108532 |
| 24 | p1 | (T)10 | 10 | 108649 | 108658 |
| 25 | p1 | (A)11 | 11 | 109134 | 109144 |
| 26 | p1 | (A)10 | 10 | 110236 | 110245 |
| 27 | p1 | (A)13 | 13 | 120967 | 120979 |
| 28 | p1 | (T)13 | 13 | 122925 | 122937 |
| 29 | p1 | (T)11 | 11 | 129082 | 129092 |

### Analysis of codon preference

Codon preference is uneven utilization of synonymous codons encoding the same amino acid in an organism (26), which is developed during the long-term evolution of organisms and has a complex set of formation mechanisms (28). The sequence analysis revealed that 87 PCGs encoded by 26,132 codons encode 87 proteins in the complete chloroplast genome of *Eupatorium fortunei* (Table 3). Among, the aliphatic group leucine appeared the most frequently with 2782 codons (10.65%), followed by isoleucine with 2201 and Serine with 1996, respectively, whereas, cysteine appeared the least frequent with 293 (1.12%). It was also observed the termination codon TAA is most common (52), which was higher than TGA (15) and TAG (20). *Eupatorium fortunei* cp has a codon preference for AT, where 70% of codons end at A/T, which is consistent with the preference of *Magnoliophyta* chloroplasts for codons ending with A/T.

**Table 3.** Codon usage in *Eupatorium fortunei*

| Codon | Amino acid | Fraction | Frequency | Number | Codon | Amino acid | Fraction | Frequency | Number |
| --- | --- | --- | --- | --- | --- | --- | --- | --- | --- |
| GCA | Ala(A) | 0.288 | 15.575 | 407 | CCA | Pro(P) | 0.289 | 12.246 | 320 |
| GCC | Ala(A) | 0.162 | 8.725 | 228 | CCC | Pro(P) | 0.179 | 7.577 | 198 |
| GCG | Ala(A) | 0.108 | 5.817 | 152 | CCG | Pro(P) | 0.151 | 6.391 | 167 |
| GCT | Ala(A) | 0.442 | 23.879 | 624 | CCT | Pro(P) | 0.382 | 16.187 | 423 |
| TGC | Cys(C) | 0.28 | 3.138 | 82 | CAA | Gln(Q) | 0.761 | 27.782 | 726 |
| TGT | Cys(C) | 0.72 | 8.074 | 211 | CAG | Gln(Q) | 0.239 | 8.725 | 228 |
| GAC | Asp(D) | 0.197 | 8.036 | 210 | AGA | Arg(R) | 0.315 | 19.019 | 497 |
| GAT | Asp(D) | 0.803 | 32.68 | 854 | AGG | Arg(R) | 0.11 | 6.62 | 173 |
| GAA | Glu(E) | 0.741 | 37.846 | 989 | CGA | Arg(R) | 0.218 | 13.126 | 343 |
| GAG | Glu(E) | 0.259 | 13.24 | 346 | CGC | Arg(R) | 0.064 | 3.865 | 101 |
| TTC | Phe(F) | 0.347 | 19.822 | 518 | CGG | Arg(R) | 0.073 | 4.401 | 115 |
| TTT | Phe(F) | 0.653 | 37.272 | 974 | CGT | Arg(R) | 0.221 | 13.317 | 348 |
| GGA | Gly(G) | 0.396 | 26.902 | 703 | AGC | Ser(S) | 0.06 | 4.592 | 120 |
| GGC | Gly(G) | 0.111 | 7.539 | 197 | AGT | Ser(S) | 0.2 | 15.269 | 399 |
| GGG | Gly(G) | 0.167 | 11.365 | 297 | TCA | Ser(S) | 0.21 | 16.072 | 420 |
| GGT | Gly(G) | 0.326 | 22.118 | 578 | TCC | Ser(S) | 0.152 | 11.595 | 303 |
| CAC | His(H) | 0.237 | 5.587 | 146 | TCG | Ser(S) | 0.078 | 5.931 | 155 |
| CAT | His(H) | 0.763 | 17.947 | 469 | TCT | Ser(S) | 0.3 | 22.922 | 599 |
| ATA | Ile(I) | 0.311 | 26.175 | 684 | ACA | Thr(T) | 0.314 | 15.613 | 408 |
| ATC | Ile(I) | 0.204 | 17.182 | 449 | ACC | Thr(T) | 0.184 | 9.146 | 239 |
| ATT | Ile(I) | 0.485 | 40.869 | 1068 | ACG | Thr(T) | 0.1 | 4.975 | 130 |
| AAA | Lys(K) | 0.727 | 38.688 | 1011 | ACT | Thr(T) | 0.402 | 20.014 | 523 |
| AAG | Lys(K) | 0.273 | 14.503 | 379 | GTA | Val(V) | 0.375 | 20.32 | 531 |
| CTA | Leu(L) | 0.14 | 14.924 | 390 | GTC | Val(V) | 0.129 | 7.003 | 183 |
| CTC | Leu(L) | 0.069 | 7.309 | 191 | GTG | Val(V) | 0.142 | 7.692 | 201 |
| CTG | Leu(L) | 0.062 | 6.582 | 172 | GTT | Val(V) | 0.354 | 19.172 | 501 |
| CTT | Leu(L) | 0.215 | 22.922 | 599 | TGG | Trp(W) | 1 | 17.488 | 457 |
| TTA | Leu(L) | 0.308 | 32.833 | 858 | TAC | Try(Y) | 0.189 | 7.041 | 184 |
| TTG | Leu(L) | 0.206 | 21.889 | 572 | TAT | Try(Y) | 0.811 | 30.308 | 792 |
| ATG | Met(M) | 1 | 24.032 | 628 | TAA | Stop | 0.598 | 1.99 | 52 |
| AAC | Asn(N) | 0.223 | 10.868 | 284 | TAG | Stop | 0.23 | 0.765 | 20 |
| AAT | Asn(N) | 0.777 | 37.923 | 991 | TGA | Stop | 0.172 | 0.574 | 15 |

### Expansion and Contraction of Border Regions

The expansion and contraction of IRs may cause changes in the SSC, an important factor for creating variation in the chloroplast genome whereas, length of its specific position and interval is an important evolutionary feature among species (26). Therefore, comparing the boundaries and adjacent genes of nine *Asteraceae* species with those of *Eupatorium fortunei*, the expansion and contraction diversification of the connected regions has been analyzed (Fig 2). However, *Asteraceae* species cp genomes are relatively conserved in terms of gene arrangement, genome structure, number of genes whereas, they all have a typical chloroplast genome structure with same boundary genes i. e., *rps19, ndhF, ycf1* and *trnH-GUG* among all species. The LSC-IRb chloroplast genome boundaries are very similar, the lengths of the LSC regions were very similar in all species except *Ageratina fastigiata*. In *Eupatorium fortunei*, the IRb-SSC boundary is *ycf1* gene whereas, IRa-SSC boundary is located within *ycf1* gene. The 49 bp length of the *ycf1* gene is located in the SSC and the remaining 569 bp is located in the IRa region, resulting formation of a pseudo-gene, different from the other nine species. The *ndhF* gene is located between 124,234 bp - 126,459 bp in the SSC region of the *Eupatorium fortunei* cp genome, with a total length of 2,225 bp, consistent with the other selected species.

**Fig 2.**
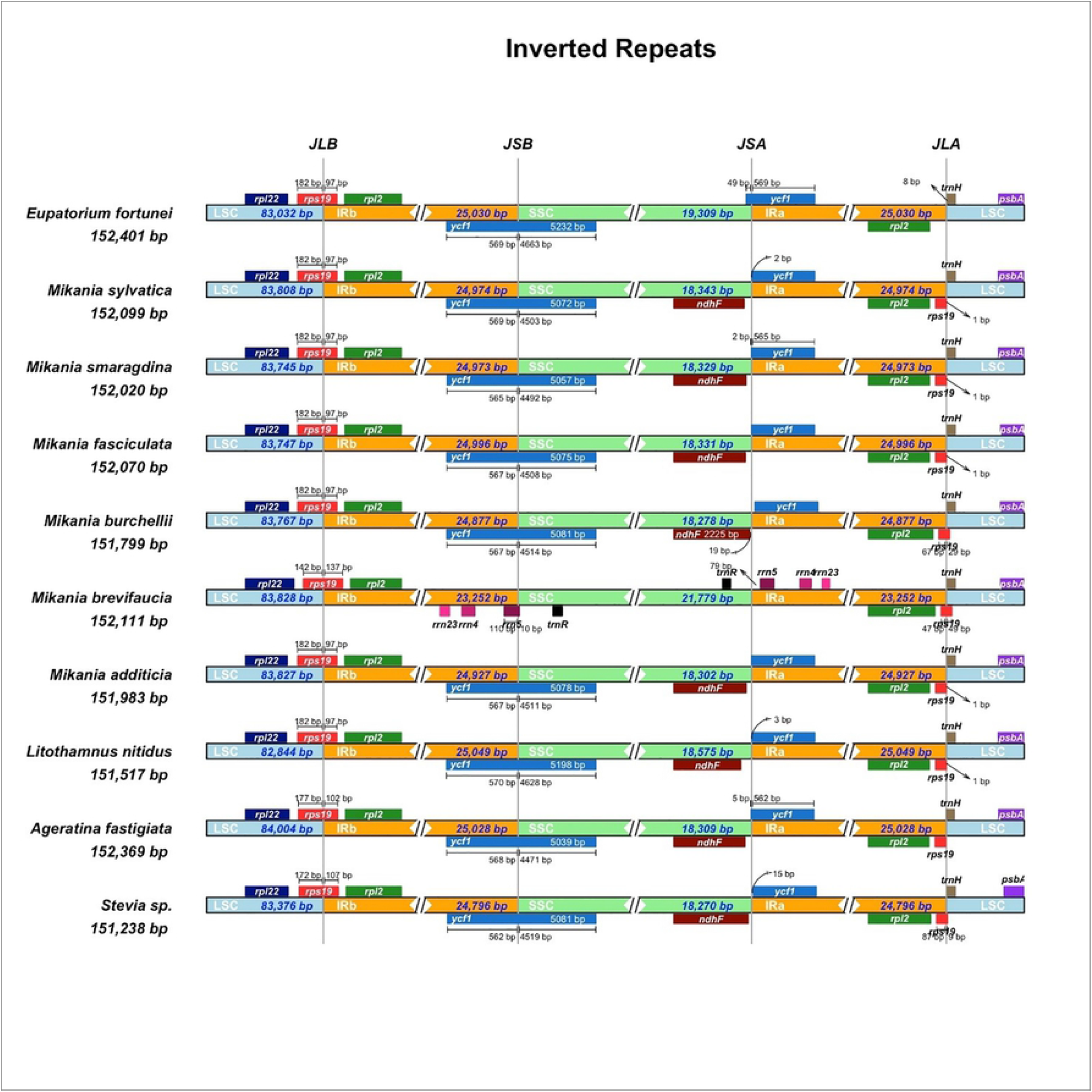
Comparison of large single copy, inverted repeat, and small single copy border regions among ten *Asteraceae* chloroplast genomes. Gene names are indicated in boxes.

In terms of length, the chloroplast of *Eupatorium fortunei* is 1.16kb which is longer than *Stevia sp*. whereas, 0.6kb, 0.4kb and 0.88kb longer than *Mikania burchellii, Mikania sylvatica* and *Litothamnus nitidus* respectively however, only 32bp longer than *Ageratina fastigiata*. Further analysis revealed that *Eupatorium fortunei* LSC region was the smaller among the nine *Asteraceae* chloroplasts whereas, the IR region was larger, suggesting *Eupatorium fortunei* IR region expanded and caused modifications along sequence length of its entire genome. Variations present in IR/SC boundary region in 10 *Asteraceae* cp genomes found responsible for differences in the lengths of the four regions and whole genome sequences.

### Sequence diversity analysis of chloroplast genomes

The sequence similarity of 13 *Asteraceae* chloroplasts was analyzed via mVISTA whole gene sequence alignment tool, which showed *Asteraceae* cp genomes showed significant sequence similarity among each other, indicating that genome structure is relatively conserved at gene sequence level (Fig 3). In particular, PCGs had a high similarity above than 95%. The gene spacer region of chloroplasts has applications for species phylogeny, molecular identification, and DNA barcoding. This study found that sequence differences between coding sequences, non-coding sequences, and spacer regions were greater than those of coding sequences. Among, *ndhD-ccsA*, *psbI-trnS*, *trnH-psbA, ndhF-ycf1* and *ndhI-ndhG* were significantly different with sequence similarity below 85%. The analysis results provided data for identification of candidate sequence loci for new *Asteraceae* plant for phylogenetic studies.

**Fig 3.**
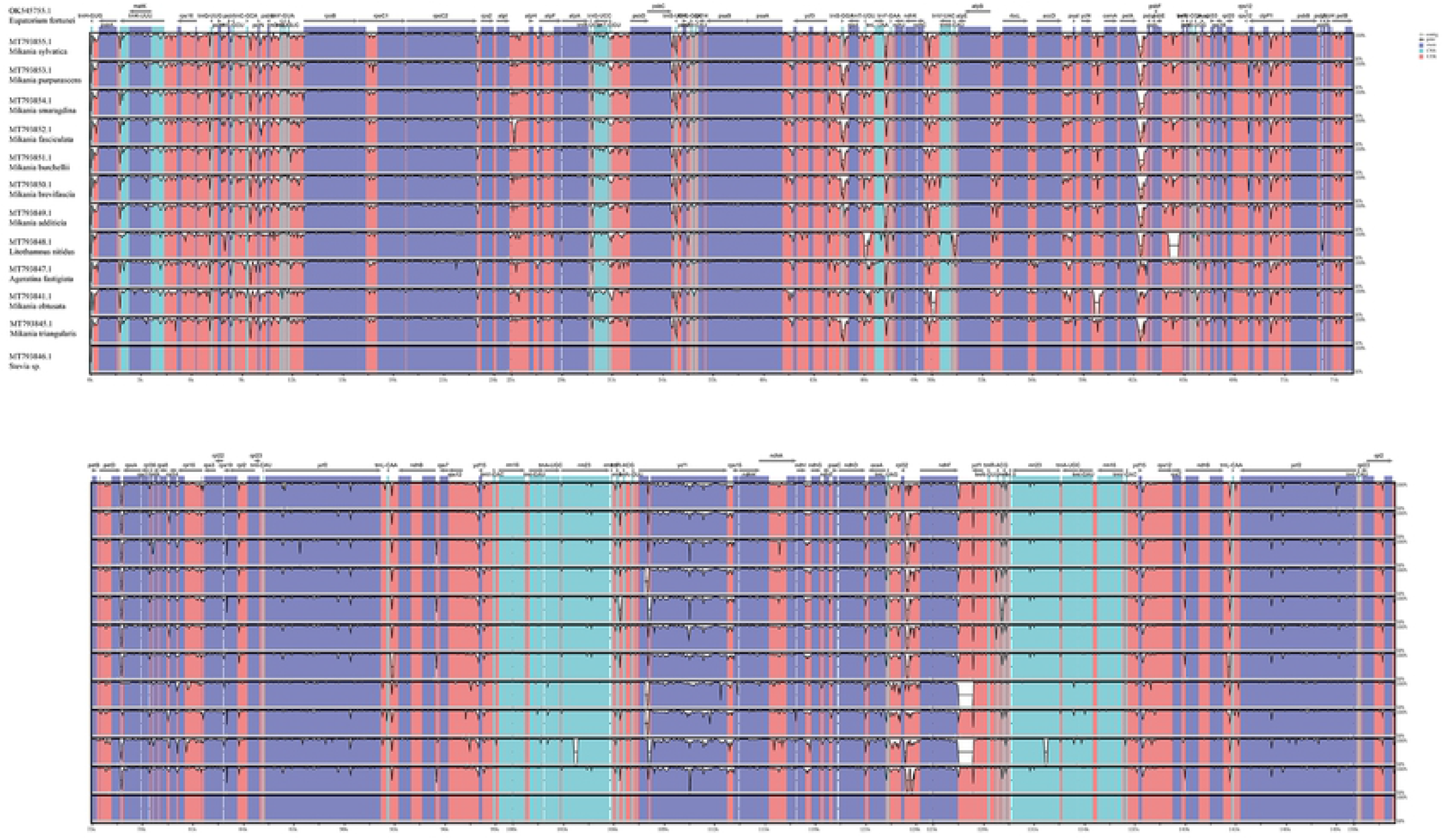
Percent identity plot for comparison of thirteen *Asteraceae* chloroplast genomes using mVISTA. Gray arrows represent genes, direction indicates forward or reverse transcription, blue represent exons, cyan represents introns, red represents spacer regions, CNS is a conserved non-coding sequence

### Phylogenetic analysis

Phylogenetic trees were generated using the complete chloroplast genome sequences of 13 species to assess the phylogenetic relationships of the *Asteraceae* plants with *Eupatorium fortunei* (Fig 4). Evolutionary relationships were inferred using the Neighbor-Joining method, offering best tree with a sum of 0.04117483 branch length. The units of branch length are the same as those used to infer the evolutionary distance of the phylogenetic tree, calculated using maximum complex likelihood method in units of base substitutions per locus. The following analysis involved 13 nucleotide sequences. All positions containing gaps and missing data were removed, resulting a total of 147,278 positions in final dataset. The resulting evolutionary tree was found to have high bootstrap support for most evolutionary branches, and phylogenetic tree was consistent with the traditional morphology-based taxonomy of *Asteraceae*, nine plants from *Mikania* forming a well-supported monophyletic evolutionary branch. In comparison, *Eupatorium fortunei* is more closely related to *Litothamnus nitidus*, a member of the same branch, with bootstrap value of 100%. As a result, it can be deduced that they are the most closely related species.

**Fig 4.**
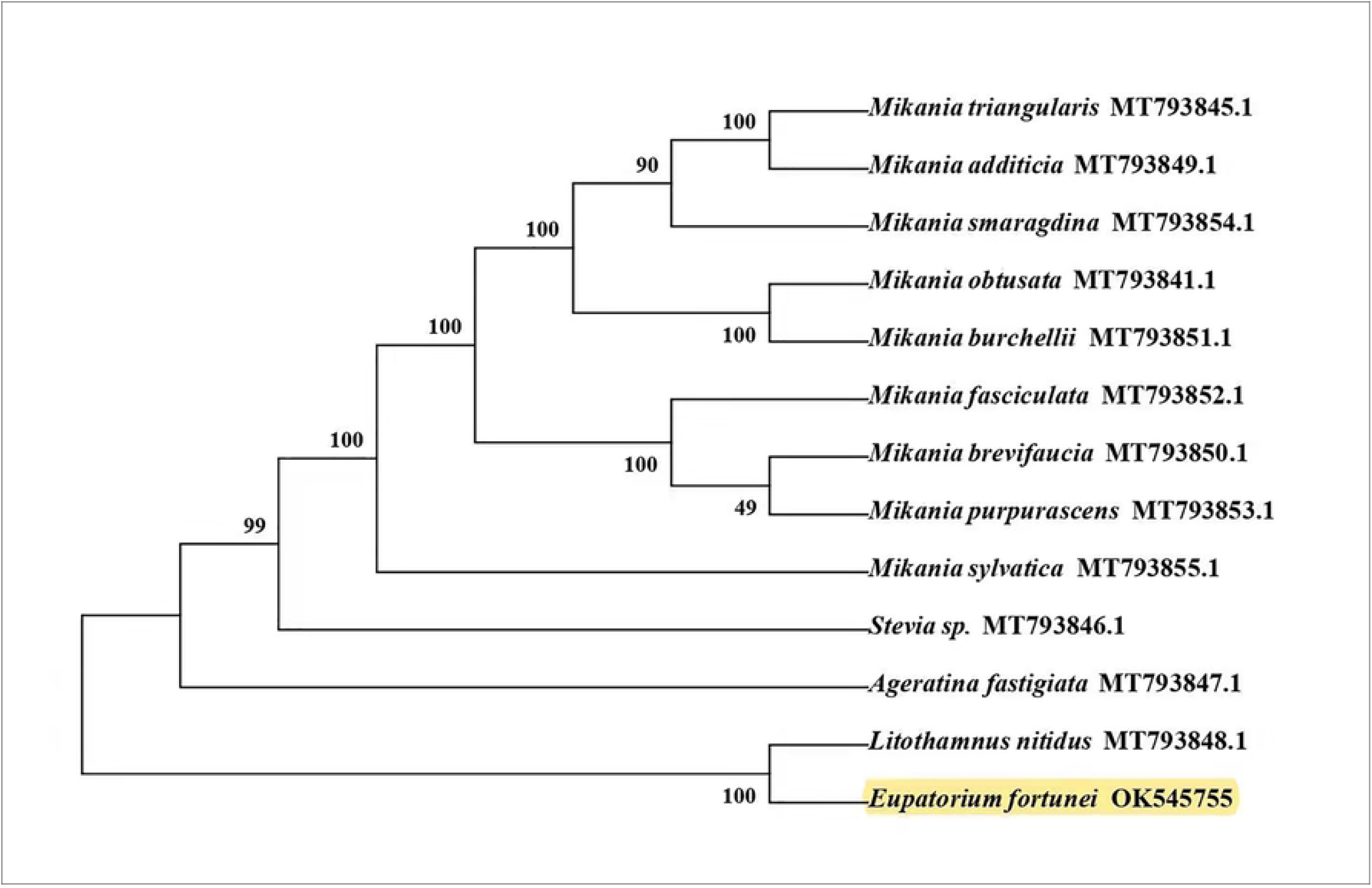
Phylogenetic tree of 13 species based on complete chloroplast genome sequences using NJ (with 1000 replicates) methods. The number below the branches indicate the corresponding bootstrap support values from the NJ trees.

## Discussion

*Asteraceae* plants are found in lands on all continents of the world, less in the tropics. In the second expression of diversity variation, studies of *Asteraceae* plants found diverse changes in inflorescence morphology and chromosome number (29). From the functional point of view, *Asteraceae* plants include important economic food crops (1), herbs (30), flower ornamental plants, and some invasive species that can have a huge impact on the ecological environment, such as *Praxelis clematidea, Ageratinn adenophora, Pityosis* et al (2, 31, 32). Because *Asteraceae* plants are rich in species, similar in phenotype, relatively late in origin, in a strong differentiation stage, and at the same time there are many intermediate links in evolution. The division of its family level and systematic research have caused great difficulties, and the systematic relationship under its family has always been a hot issue in botany research.

The highly conserved nature and low evolutionary rate of the chloroplast genome makes it a hotspot for phylogenetic studies among different species, thus making the whole sequence of the chloroplast genome a valuable tool for studying molecular phylogeny and ecology. The genetic relationships and evolutionary characteristics of medicinal plants are studied from chloroplast genome with more accuracy. In present study, we sequenced the DNA of *Eupatorium fortunei* using high-throughput sequencing platform Illumina HiSeq 6000 and obtained the complete chloroplast genome sequence of *Eupatorium fortunei* after splicing, assembly, and hole filling. The entire *Magnoliophyta* chloroplast genome length ranged from 114,914 bp to 217,942 bp. The *Eupatorium fortunei* chloroplast genome found 152,401 bp in length with 133 genes having no loss of single-copy genes, with typical *Magnoliophyta* plant chloroplast structure (28). Which is similar with the other reported *Eupatorium* plants cp genomes, The *E. catarium* cp genome is 151,410 bp in length (2). *Ageratinn adenophora* cp genome length is 150,698 bp (31).

The cp genome of *Eupatorium fortunei* has a double-stranded cyclic tetrad structure, and the junctions of SSC/IR and LSC/IR serve as evolutionary markers, which shown to be substantially conserved in size and gene capacity when compared to closely related species. The *Eupatorium fortunei* cp genome structure and gene species were highly conserved however, there were differences in genome size indicating genetic differences. The IR linker regions were compared and found to be different in the chloroplast genome among *Asteraceae* species (25). Overall, the selected species were more conserved in the LSC/IRb, IRa/LSC border regions, and in IRb/SSC, SSC The /IRa border zone is variable,, differences may be due to the contraction and expansion of the border regions. It was also observed both IRb/SSC and IRa/LSC regions were the main cause of sequence length differences in the cp genome(34), while such regions were also found in most *Magnoliophyta* chloroplast genomes.

The *Eupatorium fortunei* chloroplast genome has a codon preference for A and T, especially at second and third positions of the codon. Microsatellites can be divided into mono-, di-, tri-, tetra-, penta-, and hexa-nucleotide repeats. The locations of SSRs have functional roles in the genome, including gene regulation and evolution. In *Eupatorium fortunei* cp genome SSRs are mainly found in mononucleotide repeats, located at the LSC region. Identifying SSRs will help to advance population genetics studies, and microsatellite markers have became a powerful tool for measuring population genetic diversity and solving genetic problems, gene origins and both intraspecific and interspecific variation. Genome annotation of the whole chloroplast sequences of *Asteraceae* plants using VISTA software revealed that coding regions were more conserved than non-coding regions.

Based on the correlation of all cp genomes, the taxonomic position and evolutionary relationship of *Eupatorium fortunei* was revealed by comparison with a variety of *Asteraceae* plants. In present investigation, it was observed *Eupatorium fortunei* was most closely related to *Litothamnus nitidus*, followed by *Ageratina fastigiata* and *Stevia* sp, respectively.. Comparative analysis at genome-wide level was performed in conjunction with 12 other *Asteraceae* chloroplasts, which will bethe third complete chloroplast genome of the genus *Eupatorium* to be published on NCBI, providing genetic resources for the development of *Eupatorium fortunei* chloroplast based molecular markers, which could be the useful information for the phylogeny and molecular evolution of *Eupatorium* and even *Asteraceae*.

## Data Availability Statement

The data that support the findings of this study are openly available in GenBank of NCBI at https://www.ncbi.nlm.nih.gov/nuccore/OK545755. The associated number is OK545755.

## Acknowledgments

This study was funded by the Lanzhou Talent Innovation and Entrepreneurship Project (2017-RC-39), Gansu Provincial Science and Technology Program (20JR10RA225), Lanzhou Jiaotong University Youth Science Fund (2017007), and Lanzhou Jiaotong University-Tianjin University Innovation Fund (2018067), and we thank Lei Chen of Shanghai Yuanxin Biomedical Technology Co.

